# Spatiophylogenetic modelling of extinction risk reveals evolutionary distinctiveness and brief flowering period as risk factors in a diverse hotspot plant genus

**DOI:** 10.1101/496547

**Authors:** Russell Dinnage, Alex Skeels, Marcel Cardillo

## Abstract

Comparative models used to predict species threat status often combine variables measured at the species level with spatial variables, causing multiple statistical challenges, including phylogenetic and spatial non-independence. We present a novel bayesian approach for modelling threat status that simultaneously deals with both forms of non-independence and estimates their relative contribution, and we apply the approach to modelling threat status in the Australian plant genus *Hakea*. We find that after phylogenetic and spatial effects are accounted for, species with greater evolutionary distinctiveness and a shorter annual flowering period are more likely to be threatened. The model allows us to combine information on evolutionary history, species biology, and spatial data, to calculate latent extinction risk (potential for non-threatened species to become threatened), and estimate the most important drivers of risk for individual species. This could be of value for proactive conservation decision-making that targets species of concern before they become threatened.

## Introduction

One of the most important tools for conservation planning and prioritization is the assessment of species threat status, allowing species to be ranked by their expected risk of extinction. Unfortunately, for some large taxa (e.g. angiosperms) less than 10% of described species have been evaluated for threat status, and even in more fully evaluated taxa, many species are Data Deficient, lacking the data required for listing (IUCN 2018). Furthermore, for many species current threat status may not reflect potential future vulnerability. This is because a species’ sensitivity to human impacts is determined by the way its biology interacts with external threatening processes such as habitat loss (Fisher *et al*. 2003; Cardillo *et al*. 2004, 2005, 2008; Fréville *et al*. 2007). This means that many species currently listed as Least Concern in the IUCN Red List have biological traits that could push them rapidly to higher threat status, if they are exposed to elevated human impacts in the future: they have high latent extinction risk (Cardillo *et al*. 2006). The ability to identify species with high latent risk by predicting inherent vulnerability could be important in forward planning to minimize future biodiversity loss as environmental change continues rapidly.

We know that the current threat status of species is not only the outcome of the rapid environmental changes to which they are exposed, but is strongly mediated by the species’ biological and ecological traits, and by the environmental context (e.g. climate) in which species live. Comparative methods have been used to model the relative and interacting effects of these different kinds of factors on threat status, with the aim of understanding the causes of species declines, inferring expected threat status for unevaluated or data-deficient species, or predicting changes in threat status as impacts intensify (Fisher & Owens 2004; Purvis *et al*. 2005; Purvis 2008; Cardillo & Meijaard 2012). In plants, analyses of extinction risk have revealed many possible biological predictors, including length of annual flowering period, pollination mode, height and growth habit (Murray *et al*. 2002, 2014; Sjostrom & Gross 2006; Fréville *et al*. 2007; Sodhi *et al*. 2008; Godefroid 2014; Leão *et al*. 2014; Cardillo & Skeels 2016). However, few strong generalities have emerged from trait-based comparative analyses of extinction risk in plants, and predictive power remains relatively low (Murray *et al*. 2002).

In addition to biological traits, geographic variables and threatening processes, it has been suggested that phylogenetic properties of lineages to which species belong might be indicators of present-day risk of extinction. Of particular interest is the possibility that evolutionary isolation, distinctiveness or age should predict species’ vulnerability to human impact. Isolation or distinctiveness can be defined in various ways and are captured by a range of metrics, but Redding *et al*. (2014) point out that these concepts can be distilled to two key phylogenetic properties, “originality” (average distance of a species to all other species in the group) and “uniqueness” (a species’ distance to its nearest relative). Many authors use the term “species age” to mean the same thing as uniqueness: time since divergence from its closest known relative. The age of higher taxa (typically crown or stem age from phylogenies) has also been considered as a predictor of the prevalence of currently-threatened species. A growing list of studies support a connection between higher extinction risk and greater taxon or species age, originality or uniqueness, across a range of taxa (Gaston & Blackburn 1997; Johnson *et al*. 2002; Meijaard *et al*. 2008; Vamosi & Wilson 2008; Redding *et al*. 2010; Verde Arregoitia *et al*. 2013). These positive associations are often interpreted in terms of long-standing theories such as taxon cycles in which taxa undergo predictable trajectories of range size or ecological specialization and fragmentation (Willis 1922; Wilson 1961; Ricklefs & Bermingham 2002), the evolution of specialist adaptations leading a lineage to an evolutionary “dead-end”with inability to re-adapt to new environments or niches (Bromham *et al*. 2016), or a tendency for older species to accumulate specialist predators or parasites (Ricklefs & Bermingham 2002).

However, opposing predictions can also be made, and are supported by some empirical evidence. Recently-diverged species, particular those formed by peripatric or budding speciation, might be more vulnerable to rapid environmental change by virtue of their small initial distributions and populations. This scenario seems to be supported in the flora of South Africa’s Cape Province, where the most threatened species are clustered on short phylogenetic branches (Davies *et al*. 2011). Furthermore, the appearance of greater originality or uniqueness of a species on a phylogeny can result from a high extinction rate in the lineage to which it belongs (Gaston & Blackburn 1997; Bromham *et al*. 2016). For such species, this might predict either elevated vulnerability (because they share traits causing extinction-proneness with close relatives that have succumbed to extinction), or elevated robustness (because they are the only species among their close relatives to have withstood extinction).

Here we infer the inherent vulnerability and latent extinction risk of species from a combination of biological traits, geographic variables, and evolutionary distinctiveness, using a new Bayesian spatiophylogenetic modelling approach. Because the residuals of comparative extinction risk models frequently show phylogenetic signal, comparative studies routinely apply methods that account for phylogenetic non-independence in the observations. On the other hand, the issue of spatial non-independence has received far less attention in extinction risk studies, despite the fact that many geographic variables and threatening processes are spatially autocorrelated (Safi & Pettorelli 2010; Jetz & Freckleton 2015; Cardillo & Skeels 2016). An analytical challenge with combining both kinds of non-independence into a single model is that threat status and biological predictors are typically measured as species-level properties, whereas spatial predictors vary continuously within species distributions and are shared among species with overlapping distributions. Previous studies have skimmed over this issue by summarizing spatial variables to a single point estimate for each species (e.g. Freckleton & Jetz 2009; Jetz & Freckleton 2015; Cardillo & Skeels 2016) but this approach unavoidably discards most of the (potentially informative) variation in spatial variables. In particular, assuming covariance decays with the distance between median or mean coordinates of the species can be misleading when species ranges vary considerably in their size and shape, losing information about the extent of overlap of different species.

Our method utilizes the full distribution of observations of spatial variables for each species, combineing these with species-level estimates of biological traits and threat status, while simultaneously accounting for phylogenetic and spatial autocorrelation in model residuals. As an empirical case study we apply the approach to *Hakea*, a diverse and well-studied plant genus with 152 species distributed widely across Australia, for which we have a near-complete, well-resolved and strongly supported phylogeny and high-resolution spatial occurrence data. Like many Australian plant genera, *Hakea* are particularly diverse in the Southwest Australian biodiversity hotspot, a region that has already suffered widespread habitat loss and is projected to undergo severe climate deterioration in coming decades.

## Methods

### Spatial, biological and phylogenetic data

Occurrence records for *Hakea* (46,730 records) were from the Atlas of Living Australia (www.ala.com) filtered of records with imprecise or doubtful spatial coordinates or uncertain nomenclature (Skeels and Cardillo 2017). Spatial data for two key climatic variables (mean annual rainfall and mean annual temperature) at 0.01 degree resolution were obtained from WorldClim (Fick & Hijmans 2017). Species-level values of height, fire-response strategy (reseeding / resprouting), and flowering period (months per year in flower) were from the Flora of Australia (Australian Biological Resources Study 2018). Range area was estimated using a convex hull drawn around occurrence points for each species, log transformed to correct skewness. Threat status for every species was coded as a binary variable (threatened / not threatened) based on the species classification under the Australian Environmental Protection and Biodiversity Conservation Act or state-level threat classification schemes for Victoria, South Australia, and Western Australia, all of which use criteria similar to the Red List. If a species appeared on at least one of the four lists, it was classified as “threatened”.

The *Hakea* phylogeny includes 137 of the 152 species, and was constructed from phylogenomic data using coalescent-based species tree methods (Cardillo *et al*. 2017). From the phylogeny we calculated the evolutionary distinctiveness (ED) of each species using the “fair proportion” (FP) metric (Isaac *et al*. 2007), implemented with the “evol.distinct” function in the R library picante (Kembel *et al*. 2010). This metric calculates a weighted sum of branch lengths along the path from the root to each tip, with the length of each branch inversely weighted by the total number of tips descended from it. This means that for an ultrametric phylogeny the total amount of evolutionary time considered is equal for all species, and higher ED scores result from a species sharing its root-to-tip path with fewer co-descendants. For comparison, we also calculated ED using the “equal splits” metric, which is similar to FP but downweights values for deeper branches (Redding *et al*. 2014). Results were nearly identical (Supplementary Figure S1), so we only report results using FP in the main text.

There were 15 species with threat status data but not in the phylogeny, and these were added to the phylogeny in the following way. Each missing species was grafted onto a randomly-selected branch within the species-group to which it belongs, based on the classification scheme of Barker *et al*. (1999). This was done 200 times to generate a set of alternative trees. We tested sensitivity of results to the placement of missing species by running the statistical model (described below) on all 200 phylogenies (see Supplementary Information). For ED, species missing from the phylogeny were assigned a missing value, because the phylogenetic placement influences the ED of a given species, and their ED values were not considered in the linear predictor of the model. However, phylogenetic placements for missing species were used in the phylogenetic random effect (described below), because they still contribute useful information to help control for phylogenetic autocorrelation in the model.

As an additional spatial predictor variable, we estimated degree of habitat loss across Australia using satellite-based land use classification models (see Dinnage and Cardillo ms).

### Statistical Model Structure

Our statistical model of threat status was implemented using a Bayesian approach in the R package INLA, which uses an integrated nested Laplace approximation to estimate the joint posterior distribution of model parameters (Rue *et al*. 2009; Martins *et al*. 2013). This provides a method for incorporating fixed and random effect predictors that vary spatially with a response variable (threat status) measured at the species level. This is achieved by “integrating” the environmental fixed effects and a spatial random field across a species’ distribution, so each species contributes a single datapoint to the likelihood, even though species have multiple occurrence records and the number of records varies among species (Figure 1a gives a conceptual illustration of the model). To model the spatial effect across the entire landscape we use a “spatial mesh”, as follows. INLA uses a stochastic partial differential equation (Lindgren *et al*. 2011) approximation to a Matérn spatial covariance function (Rasmussen & Williams 2006) to estimate a random field at mesh points across a landscape. Mesh points do not necessarily need to correspond to sampled occurrence points. Random field values at observed locations in the data are interpolated using a weighted average to the three closest points on the mesh (triangulation). In order to integrate across the distribution of each species, we average across the environmental variables and the spatial random fields of each occurrence record for a species (see Supplementary Information for details). We generated a mesh using the meshbuilder function in the INLA package. This runs a Shiny app (Chang *et al*. 2018) that allows the user to generate different meshes based on a set of parameters that can be modified interactively. We chose a mesh (Figure S1) that gave good coverage across the *Hakea* occurrence points, had good statistical diagnostics, and was large enough to avoid spatial overfitting (e.g was not too small for the choice of prior – see below). Priors were chosen based on the recommendations of Illian *et al*. (2012), and are detailed in the Supplementary Information.

**Figure 1.**
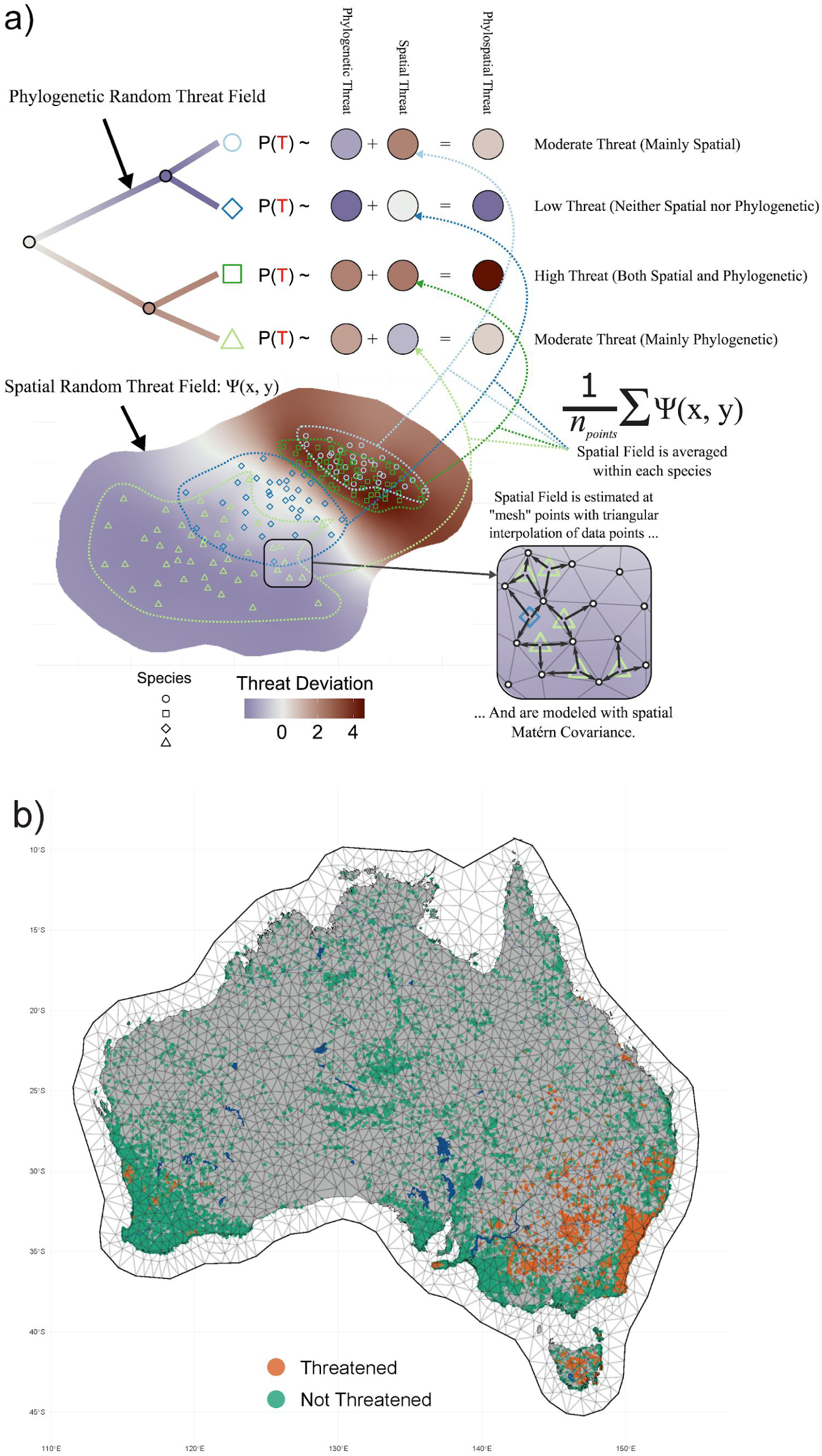
(a) Conceptual diagram of the statistical model used in this study. Here we show only the random effects, where a spatio-phylogenetic total effect is additively decomposed into a phylogenetic and a spatial effect. On the top left, phylogenetic effects are modelled as a Brownian Motion process across the phylogeny, to which spatial effect are added. The bottom half of the diagram explains the way that spatial effects are integrated across species’ full ranges. At the bottom right we show how spatial random effects are estimated across a mesh, using triangular interpolation. These spatially explicit random effects, estimated across the entire landscape, are averaged within each species to predict threat status (as indicated by dotted lines). Full mathematical details of the model can be found in the Supplementary Information. (b) The spatial mesh used in this study, with 46,730 occurrence points for 152 Hakea species listed as Threatened and non-Threatened shown as points.

INLA models covariance in a specified random effect using a precision matrix (the inverse of a covariance matrix). To model phylogenetic covariance, therefore, we used the inverse of the covariance matrix calculated from the *Hakea* phylogeny. INLA scales the precision matrix by a parameter *θ*_phylo_, which can be related to the more familiar *σ*^2^ parameter of a brownian motion covariance model (Grafen 1989) as *θ*_phylo_ = 1/*σ*^2^. The phylogenetic covariance matrix was standardised by dividing by its determinant raised to the power of 1/*N*_species_, before inversion. All predictor variables were standardised before analysis by subtracting the mean and dividing by the standard deviation.

### Comparing phylogenetic vs spatial effects

The phylogenetic and spatial random effects were modelled using different approaches, making it difficult to compare their relative strengths directly. To compare them we used the following simple approach. Rather than comparing the parameters used to fit each effect (a single scaling factor for the phylogenetic effect, and two parameters describing the Matérn covariance function for the spatial effect), we calculated the standard deviation of each random effect estimated at the species-level. In other words, we calculated the predicted deviation from the average risk for each *Hakea* species according to their phylogenetic and spatial placement, and then calculated the standard deviation of these predictions across all species, in order to measure their relative between-species explanatory power.

### Decomposing extinction risk

For each *Hakea* species we used the full model to predict the probability of being classified as threatened. We then decomposed this overall risk into contributions from the main risk factors identified in the model, defined as those for which credible intervals did not substantially overlap zero. We did this by calculating the deviation of each species from the average risk predicted from each factor independently, whilst holding all other factors constant (setting them to zero in the linear predictor). This allowed us to ask whether each species is threatened because of its phylogenetic position, geographic location, geographic features within its distribution, or biological traits.

### Mapping non-spatial risk factors

By “non-spatial” risk factors, we mean all factors besides the spatial random effect. Some of these are still explicitly spatial (climate), while others are properties of species (traits, ED, the phylogenetic random effect). These have implicit spatial structure because the species they are attached to have a spatial distribution. The strength of this statistical approach is that we can disentangle these implicit spatial effects from ‘pure’ spatial variation as represented by the spatial random field. In order to get a sense of how these risk factors are distributed on the landscape, we refit a spatial random field to the predicted threat level for each species, broken down by individual factor. That is, we calculated the predicted threat for each species based on each of the major risk factors identified in the model, holding all other factors constant. We then used this predicted risk as a new response, and refit our spatial random field using the same spatial mesh as in our original model. This generated a continuous estimate across Australia of where each risk factor has the strongest influence, based on the distribution of species with these factors.

Note that this procedure is for visualisation purposes only, not a formal statistical result, for two reasons. Firstly, a Gaussian error structure was assumed for simplicity, and while this may be a good assumption for some predicted threat values, it may not be for others. Additionally, this procedure assumes predicted threat probabilities are fixed, so does not properly propagate uncertainty in the predictions. These maps are presented in Figure 6.

### Identifying species “Of Concern”

We identified *Hakea* species with high levels of latent extinction risk based on the predicted probability of being threatened under our model, as these species may be important to consider in planning for anticipated future biodiversity loss. We classified species as being “Of Concern” if they had a predicted probability of being threatened >0.2, but were not already classified as threatened. The probability cutoff of 0.2 was chosen because it was substantially higher than the average risk predicted by the model (~0.129), and most of the species already classified as threatened had predicted probabilities higher than 0.2.

## Results

Our results did not substantially vary across the 200 randomly generated phylogenetic placements for missing species (Supplementary Figure S2), so all results presented in the main text are based on a single randomly chosen phylogeny. The effective sample size of the model estimated by INLA was 152.053, almost identical to the number of *Hakea* species in the model (152), suggesting that the approach effectively integrated the 46,730 occurrence points without inflating the statistical degrees of freedom.

The independent effects of space (Figure 2a) and phylogeny (Figure 2b) account for non-trivial amounts of variation among species in the probability of being threatened (Figure 3), though the spatial effect was roughly twice that of the phylogenetic effect. Estimated parameters for the spatial random effect suggested a spatial range of covariance with a marginal posterior mean of 25.85 decimal degrees (95% credible interval: [2.07, 121.4]), indicating a fairly large scale pattern (covariance decays to very low values only after ~25 degrees of latitude/longitude). Figure 2a shows what this spatial covariance in extinction risk looks like across Australia, and seems to suggest an east to west gradient from high to low threat. Figure 2b plots the phylogenetic random effect estimates on the *Hakea* phylogeny, revealing two major clades with higher than average extinction risk, and a large clade with relatively lower extinction risk.

**Figure 2.**
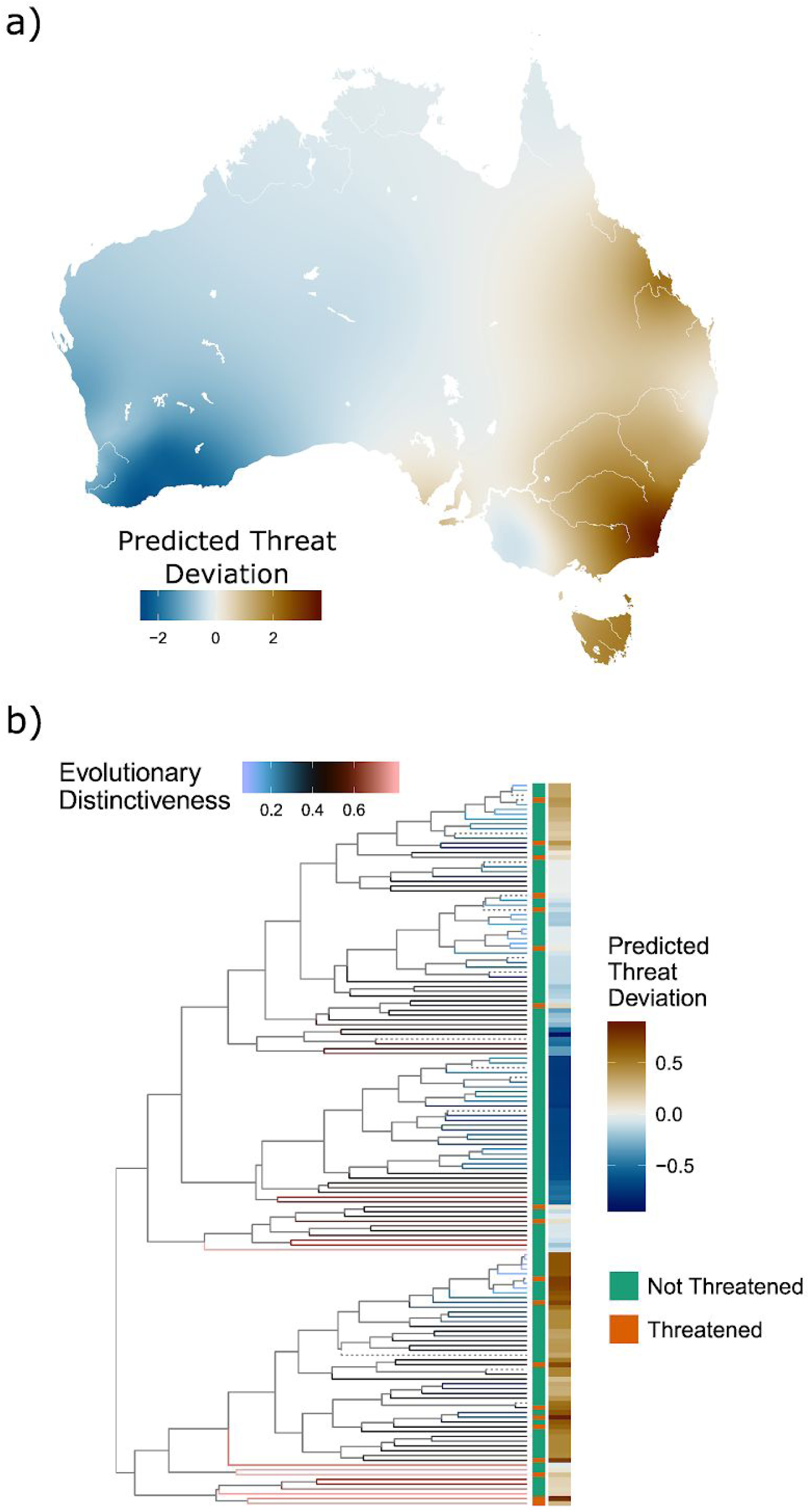
a) Map of the spatial random effect in extinction risk. Threat deviation is the predicted deviation from the average Threat probability while holding all other factors in the model constant (set to zero in the linear predictor). Predictions are made on a grid across Australia based on the fitted Matern covariance model, where values for grid points are determined by interpolation between the three closest points on the spatial mesh (see Figure 1a). b) Phylogeny for 152 species of *Hakea*. Terminal branches are coloured according to the evolutionary distinctiveness of descendant species (dotted branches lead to species that were not in the phylogeny and were placed randomly into clades; see text). Also shown are current threat status of species and the predicted deviation from the average Threat probability based purely on the species’ phylogenetic placement (according to the fitted Brownian motion model).

Three of the fixed factors in the model (ED, flowering period, and log range area) had 95% credible intervals that did not overlap with zero (Figure 3), and a fourth had a 95% confidence interval that only very slightly overlapped zero (habitat loss), suggesting a significant influence on threat probability. The other factors we tested (height, fire response, temperature, rainfall) had posterior distributions with considerable overlap with zero. We therefore consider ED, flowering period, log range area, and habitat loss to be potentially important determinants of Threat probability.

**Figure 3.**
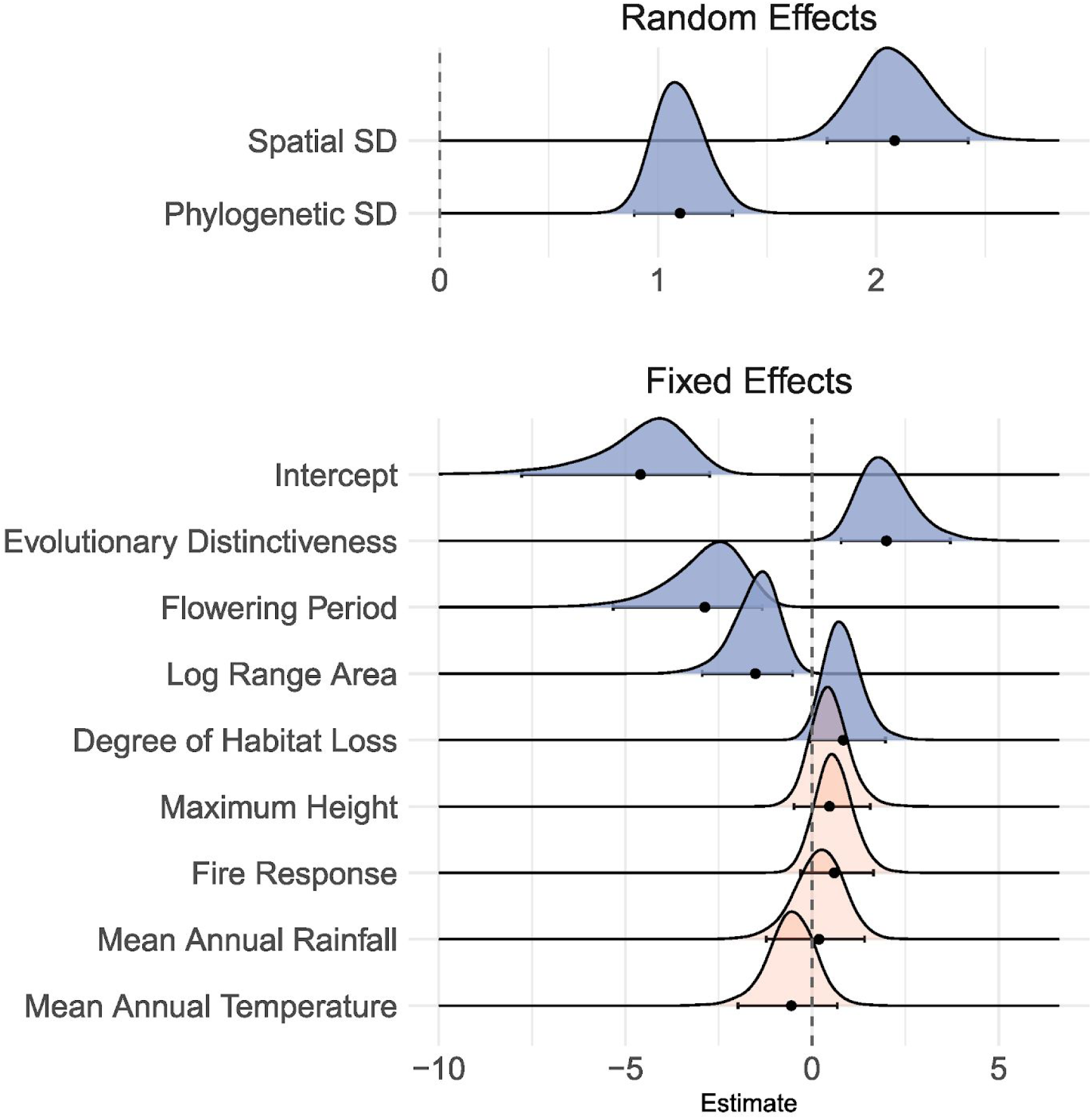
Bayesian marginal posterior distributions of model parameters. The top panel shows the estimated standard deviation (SD) of the spatial and phylogenetic random effects calculated at the species-level (see text). The bottom panel shows the linear regression coefficients of fixed effects. Because all fixed effect variables were standardised, these represent standardised coefficients, and are comparable to one another. Posterior distributions that were categorised as substantial effects, (95% credible intervals do not overlap zero) are plotted as blue; otherwise red. The mean of the posteriors are plotted as points along with error bars representing the 95% credible interval.

In general, the model clearly distinguishes *Hakea* species currently classified as Threatened from those classified as Not Threatened, with currently threatened species tending to have a predicted threat probability of >0.2, and those not currently threatened having predicted Threat probability of <0.2 (Figure 4). There are, however, a small number of currently threatened species for which the model returns a low threat probability. Conversely, there are 11 species of *Hakea* that are currently not classified as threatened for which the model returns a Threat probability of >0.2 (Figure 4, 5): these are the species with high latent extinction risk that we classified as being of concern. Figure 5b shows the geographic distribution of occurrence records for species of concern, alongside those for species currently threatened (Figure 5a). Currently threatened species are clustered into two regions of Australia heavily modified for agriculture: southeastern Australia and the Wheatbelt region of southwestern Australia. Of concern species are also clustered in these two regions, but their distribution in southeastern Australia appears to be largely around the periphery of the region occupied by currently threatened species. In southwestern Australia, of concern species are distributed across the Southwest Australian Floristic Region, including both the inland Wheatbelt zone and the high-rainfall coastal areas.

**Figure 4.**
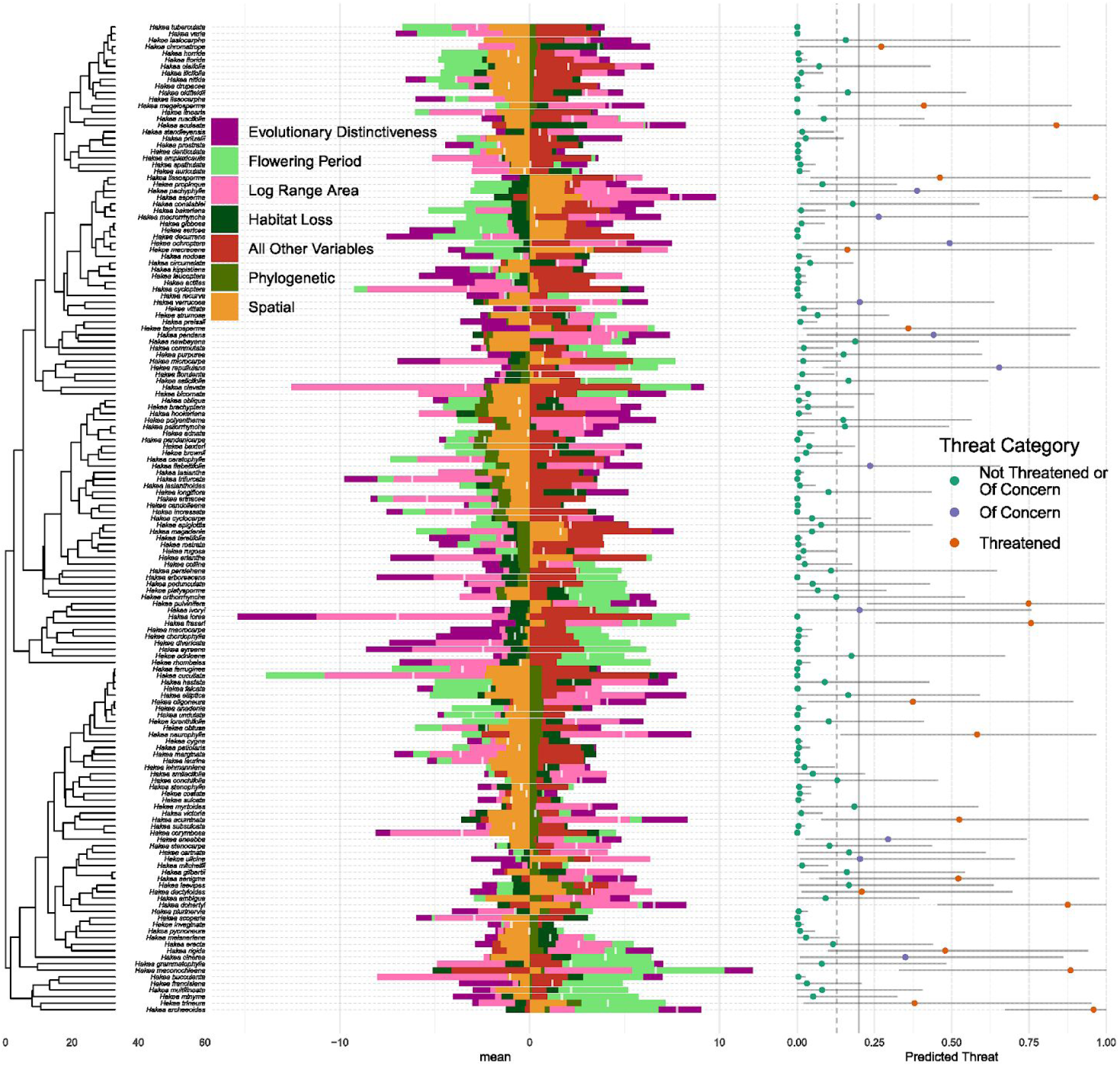
Predicted threat of all *Hakea* species from a spatiophylogenetic model, broken down into different factors determined to be substantial by the model. The panels from left to right are: 1) the *Hakea* species and their phylogeny, 2) the decomposition of the predicted threat into contributions from the different “substantial” factors and 3) the predicted probability of each species being Threatened. Predicted Threat is calculated as the predicted deviation from the average threat for each species when taking into account each risk factor independently, whilst holding other factors in the model constant (setting them to zero in the linear predictor). Error bars represent 95% credible intervals. Threat decomposition in the middle panel is split by positive and negative effects, such that stacked bars below the zero line show the relative contribution to deviations below the average, and those above the zero line show the contributions to deviations above the average. The predicted mean deviation from average threat is shown by a white bar.

**Figure 5.**
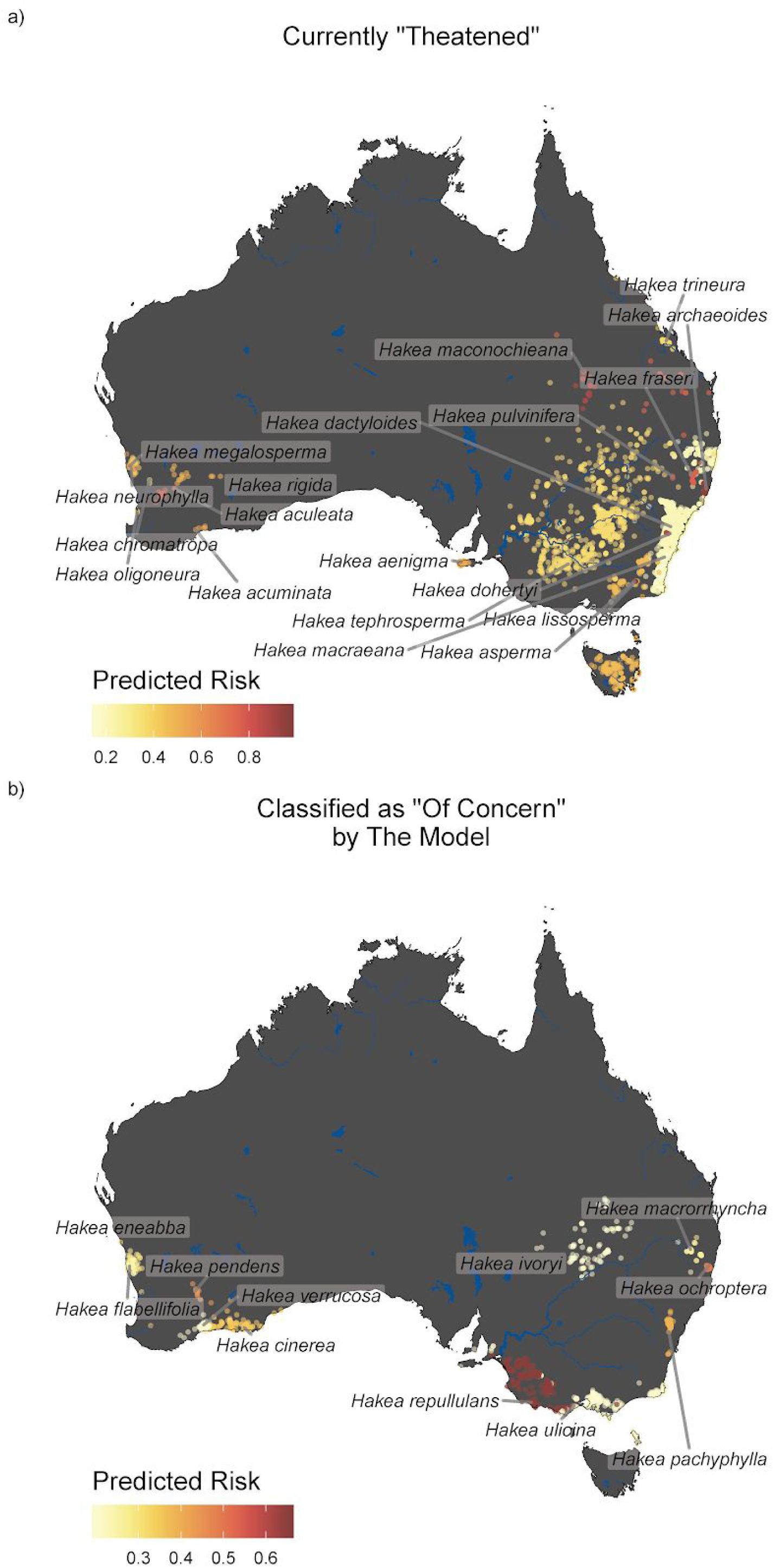
Species occurrence points for a) species currently “threatened”, and b) species the model classified as “of concern” – (not currently threatened but with predicted risk >0.20). Each occurrence point is coloured according to the species’ predicted risk.

There is substantial variation among species in the relative contributions of the two random and four fixed risk factors to the predicted Threat probability, although the influence of the fixed-effect factors tended to be larger than that of the random spatial and phylogenetic factors (Figure 4). Mapping the geographic patterns in the independent effect of each risk factor reveals a broad division between eastern and western Australia (Figure 6).

**Figure 6.**
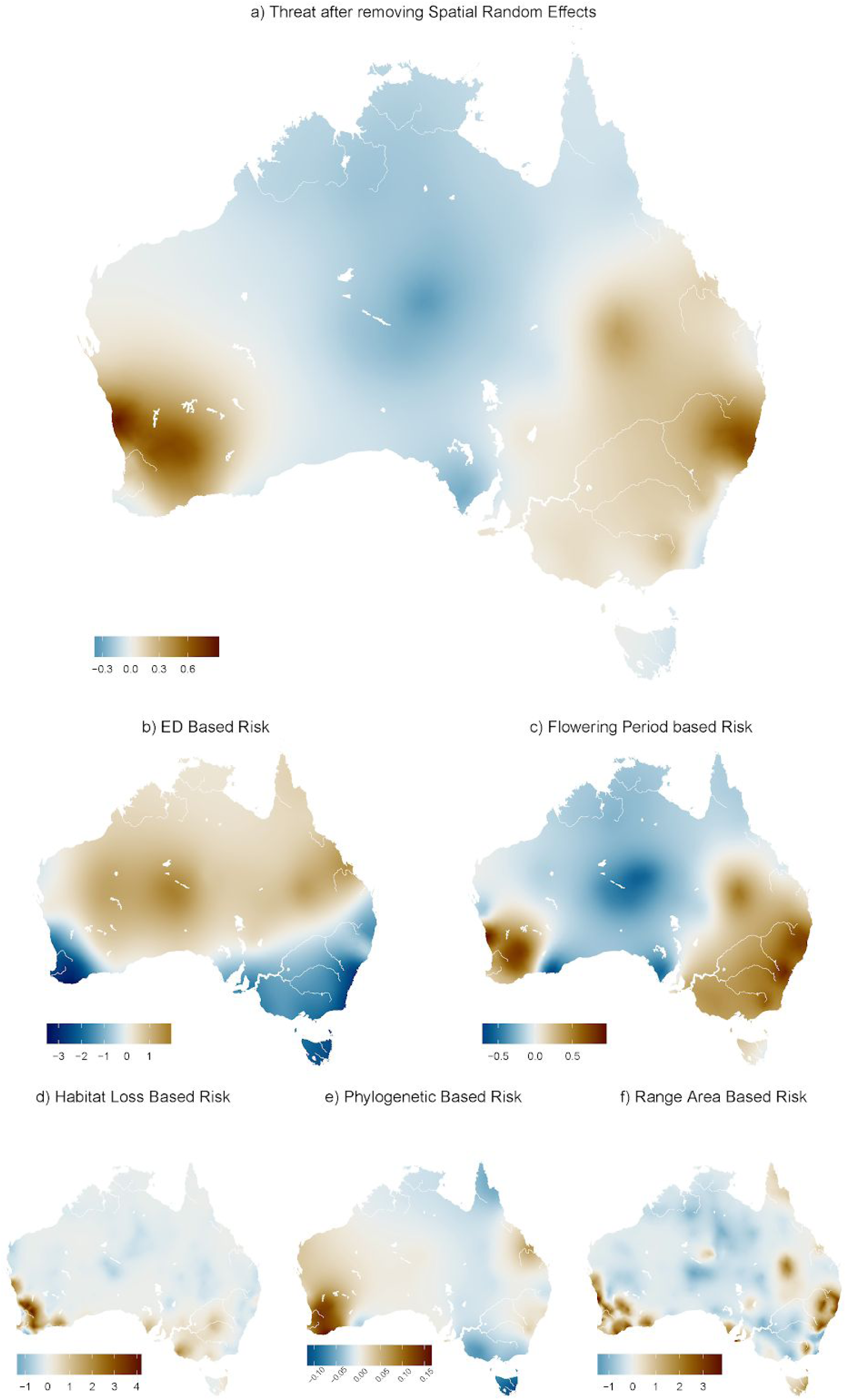
Maps of “non-spatial” risk factors. These maps plot the spatial random effects estimated on the spatial mesh for Australia (Figure 1b in main text), for the predicted risks associated with different risk factors. See methods for details on the calculations. The maps can be interpreted as a continuous approximation of how each risk factor is distributed across Australia, where risk factors associated with species traits or phylogeny is derived from the distributions of the species that they affect. For example, the phylogenetic based risk map shows where species in the most high risk clades tend to occur in Australia.

## Discussion

Our results support a link between greater evolutionary distinctiveness (ED) of *Hakea* species and a higher probability of being threatened, the first evidence for such a link that is independent of both phylogenetic and spatial effects on patterns of threat status. Although there is a growing list of taxa in which an association has been found between greater ED or evolutionary age and higher threat status (Gaston & Blackburn 1997; Johnson *et al*. 2002; Meijaard *et al*. 2008; Vamosi & Wilson 2008; Redding *et al*. 2010; Verde Arregoitia *et al*. 2013; Greenberg *et al*. 2018), the explanations for this link remain speculative. In searching for an explanation it is important to distinguish between ED, species age, and taxon age. There is longstanding evidence that taxa or clades of greater age often have a higher percentage of their currently extant species listed as threatened (Gaston & Blackburn 1997; Johnson *et al*. 2002). However, ED does not capture this aspect of evolutionary age – it is more closely connected with species age. Species age is measured from phylogenies as terminal branch length, and longer terminal branches will produce higher ED values.

If species age is driving the positive association between ED and threat status, this could be for two potential reasons (Meijaard *et al*. 2008; Verde Arregoitia *et al*. 2013; Warren *et al*. 2018, etc.): 1) Species aging effects: with increasing time since divergence from sister species, predictable changes in a species geography or biology may increase susceptibility to extinction; 2) Diversification effects: species on long terminal branches appear “old” because they belong to clades that share their high extinction vulnerability, leading to high extinction rates that leave them relatively isolated on the tree of life (“hidden extinction” effect). Alternatively, long terminal branches could result from a reduced speciation rate, and low speciation rates may be associated with elevated extinction vulnerability.

Some models of geographic range evolution suggest that species achieve their maximum range sizes at early or mid-stages of their lifespans, followed by an extended period of range contraction (Willis 1922; Jones *et al*. 2005; de Moraes Weber *et al*. 2014). Hence, the association between species age and threat status could result from smaller ranges in older species. This explanation is unlikely in *Hakea*, because there was still a positive association between threat status and ED even after accounting for range area in the model. Additionally, ED was positively correlated with range size in our study (r = 0.2; Supplementary figure S3), suggesting older *Hakea* species tend to have larger ranges. This is consistent with evidence from Paul *et al*. (2009) who showed that range sizes tend to be larger in older species of the plant genus *Psychotria*. It seems that in plants, the period of range contraction leading to final extinction might be quite short, and even if smaller range size were associated with elevated extinction risk, range size would not explain the positive association between ED and threat status.

Alternatively, other predictable biological changes, such as increasing ecological specialization with age (Meijaard *et al*. 2008), may account for the positive ED-threat association. In *Hakea*, species with narrower climatic niches tend to occupy smaller distributions, although the direction of causation is unclear (Cardillo *et al*. 2018), and again, any effect of ecological specialization on elevated extinction is unlikely to result simply from smaller range sizes. Specialization may have an effect on threat status independently of range size, but a full investigation of this effect and its relationship to species age would require data on ecologically meaningful variables (such as edaphic preferences or pollinator specificity) at relevant spatial resolutions, and such an analysis was beyond the scope of the present study.

A high ED value or a long terminal branch for a species is not necessarily an indicator of species “age”: longer branches may also result from a low rate of diversification along a lineage, either from an elevated extinction rate or a decreased speciation rate. If there are traits with phylogenetic signal that are associated with increased extinction or decreased speciation rates (see Vamosi *et al*. 2018 for a review), then a higher extinction vulnerability in contemporary species in these lineages could be the result of sharing these conserved traits. There is some evidence that some traits are associated both with extinction rates over evolutionary time and present-day threat status of species (Vamosi et al 2018). It is also possible to imagine that traits related to lower speciation rate could affect a species’ contemporary risk of extinction. For example, a clade with high rates of extinction may be characterised by a tendency to form many spatially structured sub-populations with weak gene flow between them, increasing the likelihood of speciation through isolation.

The strongest biological predictor of threat status in *Hakea* is length of flowering period – species with shorter periods in flower each year are more likely to be threatened. The same pattern has also been found in *Banksia* (Cardillo & Skeels 2016) and some other plant groups (Lahti *et al*. 1991; Fréville *et al*. 2007; Ames *et al*. 2017). Longer flowering periods could be associated with greater seed set and recruitment, and hence larger populations, which might buffer plant species against disturbances (Fréville *et al*. 2007). Longer flowering periods may also contribute to flexibility in responding to anthropogenic stresses. This could be thought of as a temporal insurance effect, analogous to the spatial insurance effect thought to underlie the negative association between range size and extinction threat (Loreau *et al*. 2003; Gaston & Fuller 2009; Keith *et al*. 2013). For example, plants with a short flowering period may be highly dependent on particular insect, bird or mammal pollinators that emerge around the same time. If these pollinators are at risk themselves, it limits the options available for reproduction. Comparatively little is known about pollinator specificity in *Hakea* – although the majority of species seem fairly flexible, some species are known to have a narrow range of preferred pollinators (Barker *et al*. 1999). Any threat processes that are temporally heterogeneous could increase the risk of interrupting reproduction for species with short flowering periods, due to this insurance effect.

The spatiophylogenetic method we have applied in this study allows us to map the geographic patterns in the independent effects of different predictors of threat status. We see from Fig. 6 that when purely spatial effects are removed, the distribution of species assemblages with high threat probability predicted from the model shows a clear geographic pattern. The two regions of highest predicted threat are centred on the central east coast and the southwest coast of Australia. This geographic pattern seems to be driven primarily by the pattern of threat predicted from flowering period alone, and to a lesser extent to the pattern predicted from phylogenetic position alone. In fact, these two predicted “threat hotspots” also overlap substantially with Australia’s two global biodiversity hotspots (southwest Australia and forests of eastern Australia; https://www.cepf.net/our-work/biodiversity-hotspots). Global biodiversity hotspots are defined on the basis of unusually high concentrations of endemic plant diversity, combined with high levels of anthropogenic land-use change (Myers *et al*. 2000). Our results suggest that for *Hakea*, hotspots are additionally regions in which species have a tendency to be “inherently” vulnerable, on the basis of a brief annual flowering period and their phylogenetic position. This coincidence of regions where (a) there is high endemic diversity, (b) a high proportion of original habitat is already gone, and (c) species are inherently vulnerable because of their biology, means that these are regions that should be afforded high priority for the protection of remaining natural habitat.

The independent effect of ED on threat status shows a largely complementary geographic pattern to those for flowering period and phylogeny (Fig 6). Here, it is the arid zone (including the inland and the south-central and west coasts) and the north of Australia which are the regions of highest predicted threat probability. This can probably be explained by the low species richness of these regions and the fact that a number of species (especially in the arid zone) are on the ends of long branches (Cardillo et al 2017), which elevates ED in these regions. One apparent consequence of the almost complementary geographic patterns for ED and the other variables is that distributions of many of the species with high “latent risk” – species not yet threatened, but predicted by the model to be threatened – lie in the zones of overlap between regions of high threat predicted from ED and from the other variables (Fig 5). These are species we have classified as being “of concern”, because their inherent vulnerability could push them rapidly towards extinction if habitat loss or other ecosystem disturbances within their distributions increase. Of particular concern are five species whose range overlaps the southwest Australian hotspot, because this is a region expected to undergo severe climate deterioration in coming decades, with projected rainfall decline of up to 35% (Council n.d.; “Climate Change in Australia” 2018).

The bayesian spatiophylogenetic method developed here solves a general problem in comparative analysis and so may be useful for addressing a variety of different questions. Many questions in comparative biology are potentially influenced by the spatial arrangement of species, yet spatial effects are still not routinely incorporated into analytical models in the way that phylogenetic effects are. This is likely due to a scarcity of detailed spatial data (at least until fairly recently) and a lack of appropriate methods, that are easily implemented, to incorporate both spatial and phylogenetic effects into comparative analyses. Our method allows both kinds of effects to be modelled simultaneously, and also offers a potential solution to another major issue for spatially explicit comparative methods, the mismatch in the levels of measurement of spatial variables and species variables. Here we have shown how the method can show spatially-explicit insights into extinction threats not possible otherwise, and allow for a more nuanced and careful exploration of comparative data on species threat status.

## Acknowledgements

Thanks go to Haakon Bakka for advice on the statistical model. This research is supported by Australian Research Council Discovery Grant DP160103942. A.S. is supported by the Australian Government Research Training Program.

## Supplementary Information

### Statistical Model Details

Mathematically, we can write out the equation of our full model as:

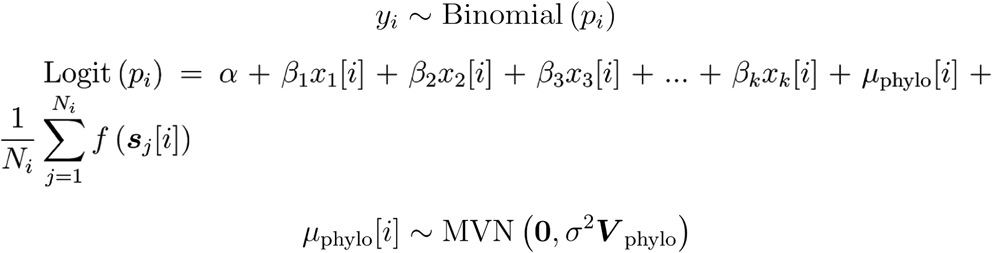

where *α* is an intercept term, *β_l_* is the *l*th fixed effect coefficient describing the effect of species-level predictor *l* of species *i* on the response, **0** is a vector of zeroes, *σ*^2^ is a phylogenetic scaling factor, ***V***_phylo_ is the standardised phylogenetic covariance matrix for all species in the model, *f*(***s****_j_*[*i*]) is a function describing the spatial effect for the *j*th longitude/latitude occurrence coordinates of species *i*: ***s****_j_*[*i*], and *N_i_* is the total number of observed occurrences of species *i*.

The spatial function can be expanded as:

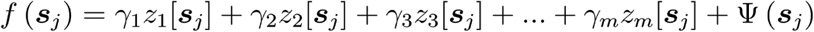

Where *γ_m_* represents a regression coefficient determining the linear effect of environmental variable *z_m_*[***s****_j_*], measured at coordinate ***s****_j_*, and Ψ (***s****_j_*) is a function describing the spatial random effect. The *γ* parameters can be moved out of *f* (***s****_j_*) because they are simple linear terms, and so the mean of their sum is the same as the sum of their means. So, if we set 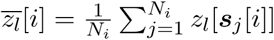, we can simplify the model to:

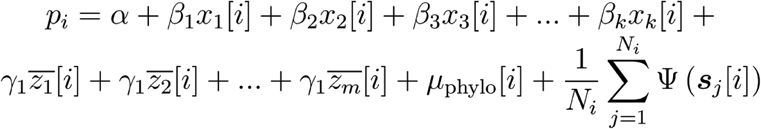

Such that the *γ* terms are now fixed effect coefficients on the mean environmental variables for each species as a whole (e.g. the mean of the variables across all of a species’ occurrence points). The Ψ (***s****_j_*) function is a basis function that approximates the spatial Matérn covariance function across a spatial mesh, using a stochastic partial differential equation approach (SPDE: Lindgren *et al*. 2011). Lindgren *et al*. (2011) has mathematical details of the SPDE approach, which we will not reproduce here.

INLA uses a set of weights to calculate the spatial random effects at the coordinates of the data. The weights interpolate between the three closest mesh points, and are encapsulated in a matrix (the ***A***_spatial_ matrix) with *N*_point_ rows (where *N*_point_ is the number of occurrence points), and *N*_mesh_ columns (where *N*_mesh_ is the number of mesh points), where the weights in each row sum to 1. In order to calculate the average spatial effect across each species’ occurrence points (e.g. 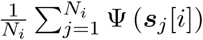) we calculated a new ***A***_species:spatial_ matrix with *N_i_* rows and *N*_mesh_ columns, where each row was simply the mean of the rows in ***A***_spatial_ corresponding to the occurrence points of each species *i*. Again, weights in all rows sum to 1 in this new ***A*** matrix.

### Choosing Bayesian priors

INLA requires priors on all parameters to be specified. For fixed parameters we used the default INLA prior, a wide gaussian prior with mean = 0 and variance = 100. For the phylogenetic and spatial random effects we used weakly informative priors, as recommended by Simpson *et al*. (2017) and Gelman *et al*. (2008). For the phylogenetic scaling parameter we used the ‘pcprior’ distribution in INLA with parameters 1 and 0.1, corresponding to an exponential distribution with about 10% of its probability distribution >1. To test the sensitivity of our analysis to the choice of prior, we ran the full model with several different prior parameters ([1, 0.01], [1, 0.1], [1, 0.5], [1, 0.99]) representing distributions with increasingly heavy tails, the last of which approximates an uninformative uniform distribution. The choice of prior had very little effect on any other parameter estimates, and all qualitative results were identical, so for all subsequent analyses we used [1, 0.1].

The priors on the spatial random effect (which include the range and *σ* parameter of the INLA implementation of the (Rasmussen & Williams 2006) were chosen as follows. Illian *et al*. (2012) recommend choosing priors on the range parameter (representing the spatial range over which the covariance decays to almost zero) that avoid spatial overfitting, by placing most of the prior density on range values greater than the apparent covariance range of the environmental factors used in the model. Allowing values much less than this can result in a spatial random effect that overfits on a very fine spatial scale, which will explain away any environmental factors, and lead to poor predictions for new data. By choosing a prior that enforces a similar spatial covariance in the spatial random effect and fixed environmental factors, we allow the model to more appropriately compare between them and choose the most parsimonious decomposition of the effects. We chose a prior through trial and error by fitting only the spatial random effect to our data, then comparing a map of the result to maps of our environmental factors until we found a set of priors where the random effect map showed a similar spatial covariance to the environmental factor maps. For the range parameter, a ‘pcprior’ with 10% of its density <2 decimal degrees resulted in appropriate covariance structure. Any value >2 in the prior resulted in very similar results that avoided overfitting (as the estimated range in the model was considerably greater than 2; see Results). On the *σ* parameter we used a ‘pcprior’ distribution with values [1, 0.01]. The *α* parameter, which controls the ‘smoothness’ of the (Matérn covariance function: Rasmussen & Williams 2006) was fixed to 2, which is a standard choice in spatial modelling.

## Supplementary Figures

**Figure S1.**
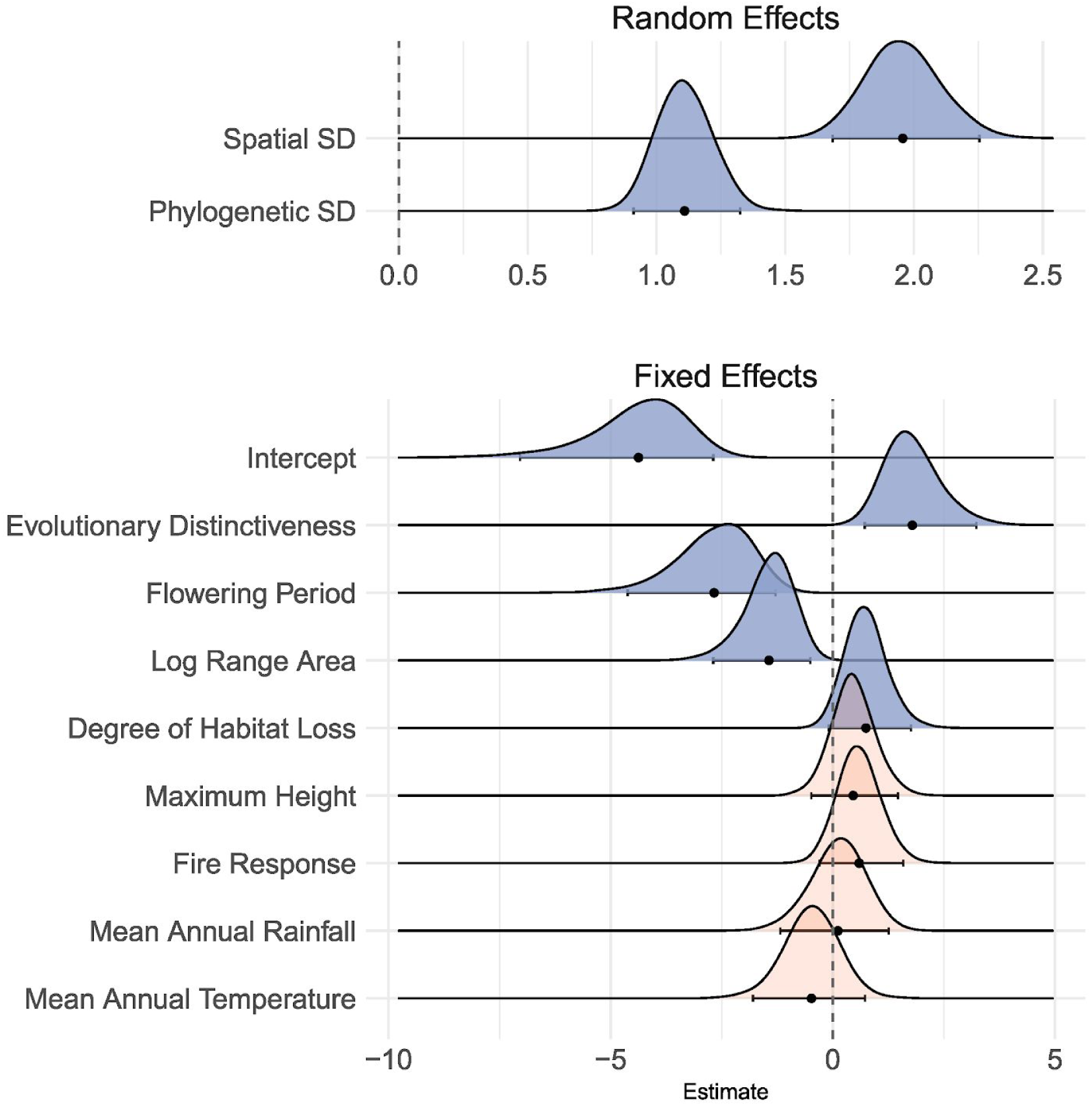
Bayesian marginal posterior distributions of model parameters when run using alternative method of calculating ED (“equal splits”). The top panel shows the estimated standard deviation of the two random effects as calculated at the species-level. The bottom panel shows the fixed effect linear regression coefficients. Because all fixed effect variables were standardised, these represent standardised coefficients, and are comparable to one another. Posterior distributions that were categorised as substantial effects, (95% credible intervals do not overlap zero) are plotted as blue; otherwise red. The mean of the posteriors are plotted as points along with errorbars representing the 95% credible interval.

**Figure S2.**
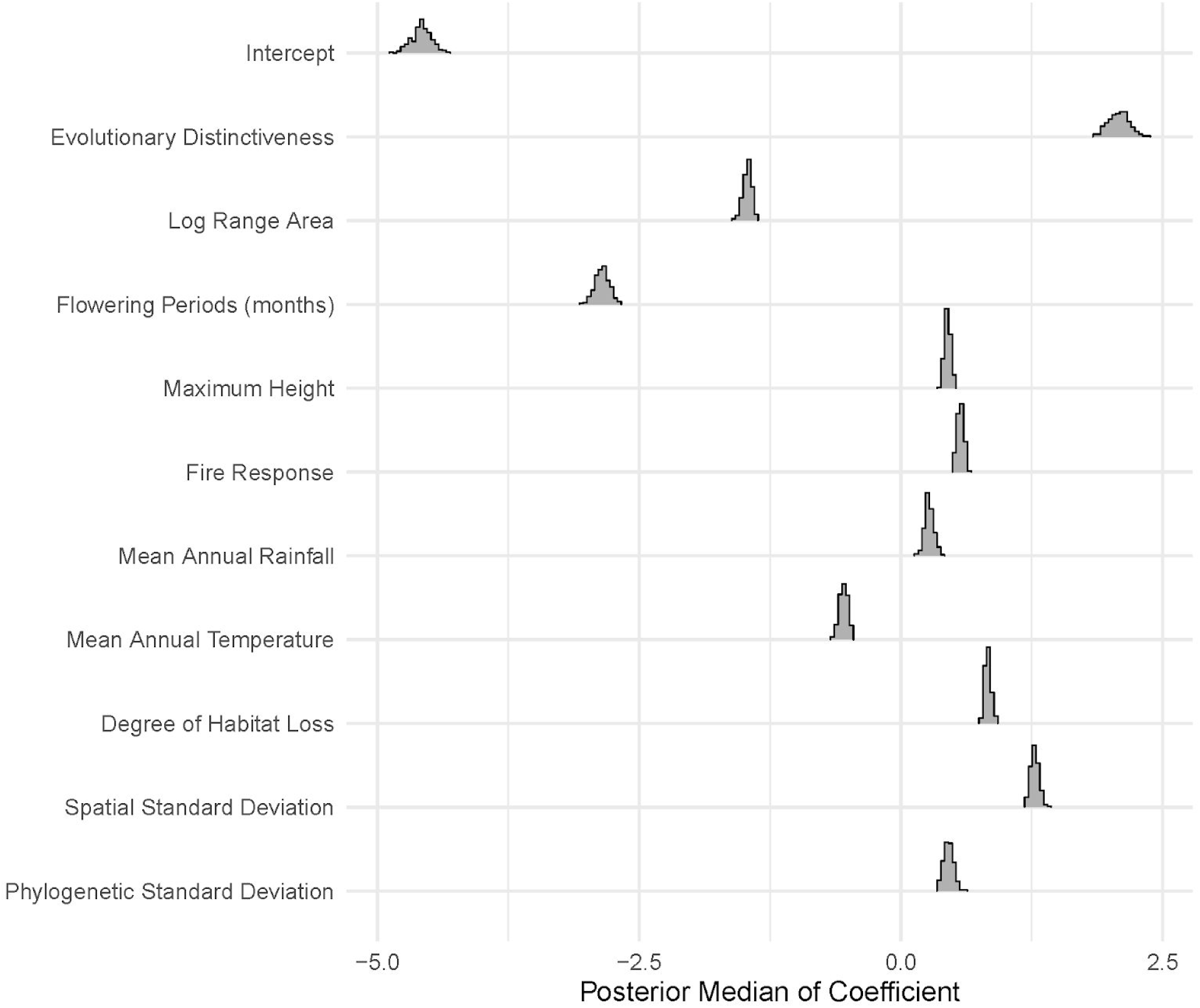
The distribution of model coefficients across models run for each of 200 generated phylogenies, where for each phylogeny all of the 15 missing species were placed randomly within a clade that corresponds to its taxonomy. Specifically, each species was placed randomly within the clade that subtends the node corresponding to the most recent common ancestor of all other species found in same taxonomic group as the missing species (according to Barker *et al*. (1999)). Coefficients were summarised for each model by the median of its marginal posterior distribution.

**Figure S3.**
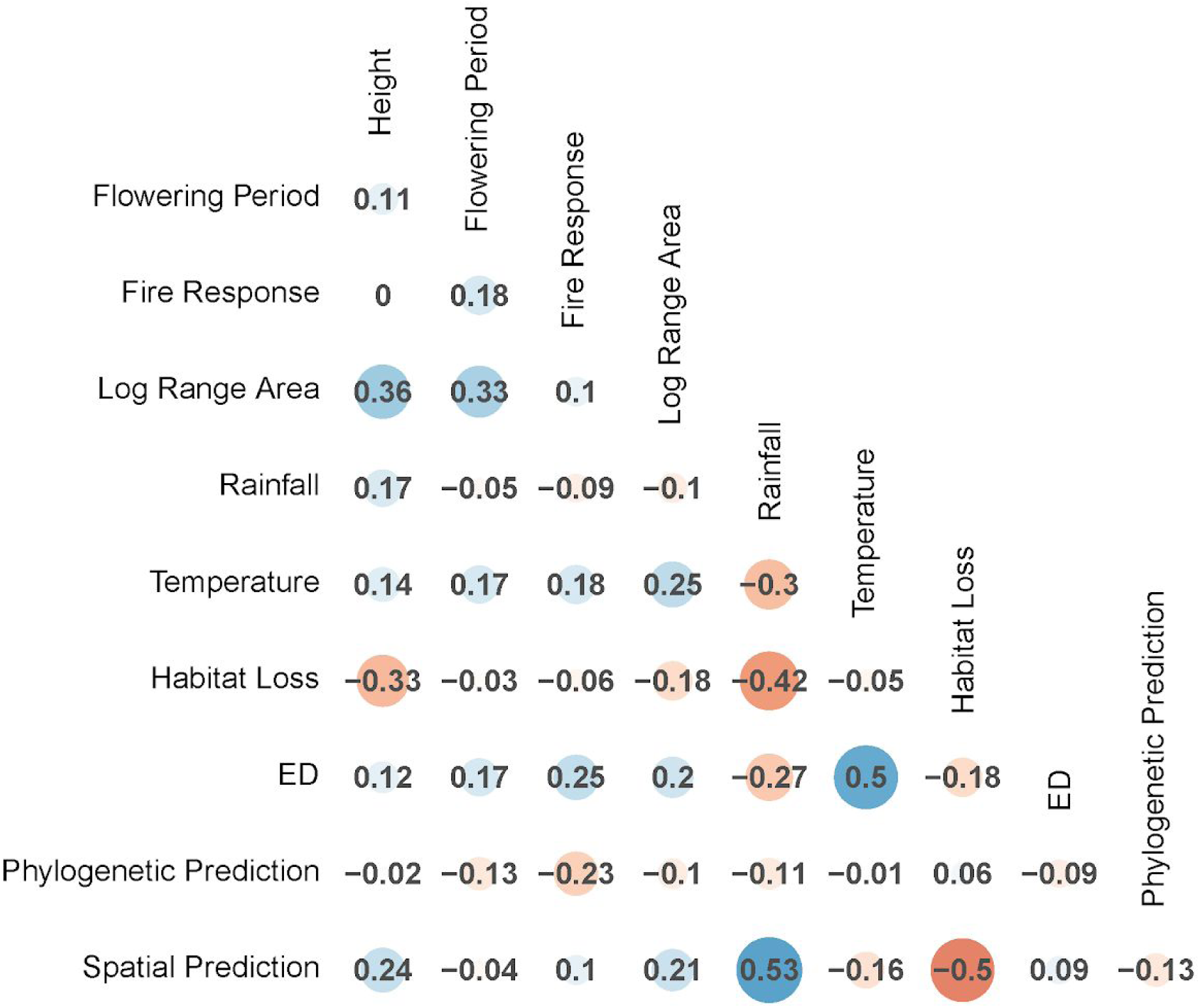
Correlation matrix of all predictors in the model. Additionally included are the phylogenetic and spatial predictions from the model.

